# Cell shape characterization, alignment and comparison using FlowShape

**DOI:** 10.1101/2022.12.08.519700

**Authors:** Casper van Bavel, Wim Thiels, Rob Jelier

**Affiliations:** Centre of Microbial and Plant Genetics, KU Leuven, Leuven, Belgium

## Abstract

**Motivation:** The shape of a cell reflects, among other things, actomyosin activity and adhesion properties. Cell shape is further tightly linked to cell differentiation and can reveal important cellular behaviors such as polarization. Hence, it is useful and informative to link cell shape to genetic and other perturbations. However, most currently used cell shape descriptors capture only simple geometric features such as volume and sphericity. We propose FlowShape, a new framework to study cell shapes in a complete and generic way.

**Results:** In our framework a cell shape is first represented as a single function on a sphere. The curvature of the shape is measured and next mapped onto a sphere in a conformal manner. This special curvature map is then approximated by a series expansion: the spherical harmonics decomposition. This decomposition facilitates a wide range of shape analyses, including shape alignment, statistical cell shape comparison and inference of cell shape deformations over time. From this representation, we can reconstruct the cell shape using the Dirac equation. The new tool is applied to perform a complete, generic analysis of cell shapes, using the early *Caenorhabditis elegans* embryo as a model case. We distinguish and characterize the cells at the seven-cell stage. Next, a filter is designed to identify protrusions on the cell shape to highlight lamellipodia in cells. Furthermore, we use our framework to identify any shape changes following a gene knockdown of the Wnt pathway. Cells are first optimally aligned using the fast Fourier transform, followed by calculating an average shape. Shape differences between conditions are next quantified and compared to an empirical distribution. Finally, we put forward a highly performant implementation of the core algorithm, as well as routines to characterize, align and compare cell shapes, through the open-source software package FlowShape.

**Availability:** The data and code needed to recreate the results are freely available at https://doi.org/10.5281/zenodo.7391185. The most recent version of the software is maintained at https://bitbucket.org/pgmsembryogenesis/flowshape/.

**Author summary:** We present FlowShape, a framework for cell shape analysis, based on the concept of *spherical harmonics* decomposition. This decomposition allows for any function defined on a sphere to be rewritten as a weighted sum of basis functions. Contrary to previous work, we use a single function to describe a shape, the mean curvature, which implies that the decomposition weights can be used as a complete shape description. The expression of a shape in this manner allows for very efficient calculations, as we illustrate with the *C. elegans* embryo as a model. The decomposition permits efficient comparison and alignment of shapes. We demonstrate this by clustering the cells in the early embryo and illustrating the different shapes by cluster. The decomposition further facilitates averaging of shapes and searching for particular features on the shape by defining filters that can then be efficiently applied. Finally, we illustrate how the framework can facilitate statistical comparisons between shapes.

## Introduction

The shape of a cell can be a window into many underlying cellular processes, since it arises from a complex interplay of cortical actomyosin contractility, directed force generation and cellular adhesion. Tightly controlled changes in shape are central to cell division, as well as cell migration and morphogenesis. [1, 2] Furthermore, cell shape reflects processes such as cell differentiation and cellular responses to signals and polarization. [3, 4]

Shape information can be leveraged for a variety of applications. For example, measuring curvatures and angles across cell contacts can inform about cortical tensions and cell-cell adhesion strengths. [5, 6] It is commonly used to identify the cell type or state [7], and can serve to identify phenotypes following a perturbation [8]. Cell shape analysis can also uncover cellular mechanical properties. These kinds of applications commonly make use of simple shape descriptors, such as sphericity and eccentricity. However, such descriptors highlight only particular features of a shape and do not fully describe one. Given that these descriptors only capture a particular facet of cell geometry, multiple descriptors are often combined to get a more complete characterization. [9]

A more direct and versatile approach would be to use a formalism that can completely and generically describe a shape. One such formalism is *spherical harmonics* (SH). SH are a set of frequency-space basis functions on the sphere. They can generically describe any cellular shape, and further provide a powerful mathematical framework to make characterizations, generalizations and comparisons between shapes in a computationally efficient manner. First, a shape is represented as a function on a sphere by a mapping procedure. Second, the functions on the sphere can be decomposed into a linear combination of SH, analogous to the Fourier series of a one-dimensional periodic function. [10]

SH have been applied successfully to describe shapes in the life sciences, for example to represent and compare brain anatomy [11] and other organs [12], as well as parts of embryonic development, such as mouse limb buds and embryonic hearts [13]. They have also been explored to describe cell shapes and dynamics, but despite some promising results, they have not attracted much attention in the field. [14–19] In these applications, shapes mapped to the sphere are represented by three coordinate functions, which limits the advantages of using SH. A broader application of the method may further have been hindered by the lack of an efficient and readily useable software package.

Here we propose FlowShape, a framework to describe cell shapes completely and to a tunable degree of detail. First, the procedure maps the mean curvature of the shape onto the sphere, resulting in a single function. Next, this function is decomposed into SH to capture shape information. This SH representation is then used to align, average and compare cell shapes, as well as to detect specific features, such as protrusions. The package is written in Python and is freely available. To evaluate and illustrate our approach, we apply it here on the cells of the early *C. elegans* embryo.

### Design and implementation

FlowShape works directly on discrete triangle meshes that represent shapes. These meshes were reconstructed from confocal fluorescent microscopy imaging of live *C. elegans* embryos which express a membrane-tagged fluorescent protein. The resulting images were segmented using the SDT-PICS procedure [20], which outputs a closed triangle mesh representing the cell membranes at each time point (fig. 1A). We then use SH as shape descriptors to comprehensively describe the shape. To describe these cell shapes using SH, we first need to map the shape to a function on a sphere, which can be done efficiently by mean curvature flow. Intuitively, mean curvature flow repeatedly applies a smoothing operation until the surface converges to a sphere. As mean curvature flow can develop singularities, we use the conformalized mean curvature flow, which avoids numerical instabilities and is completely stable. [21] Once a map is found, the surface should be represented on the sphere by one or more functions. Here a single function is used, the mean curvature. This function is sufficient to represent the shape if the map to the sphere is *conformal*, i.e. it does not distort angles (fig. 1B). The conformalized mean curvature flow fulfills this requirement. SH can give a compact representation of any function on a sphere, as a finite set of coefficients. To obtain the coefficients that weigh the contribution of each SH, a least squares optimization is performed, using the iterative residual fitting method. [22]

**Fig 1.**
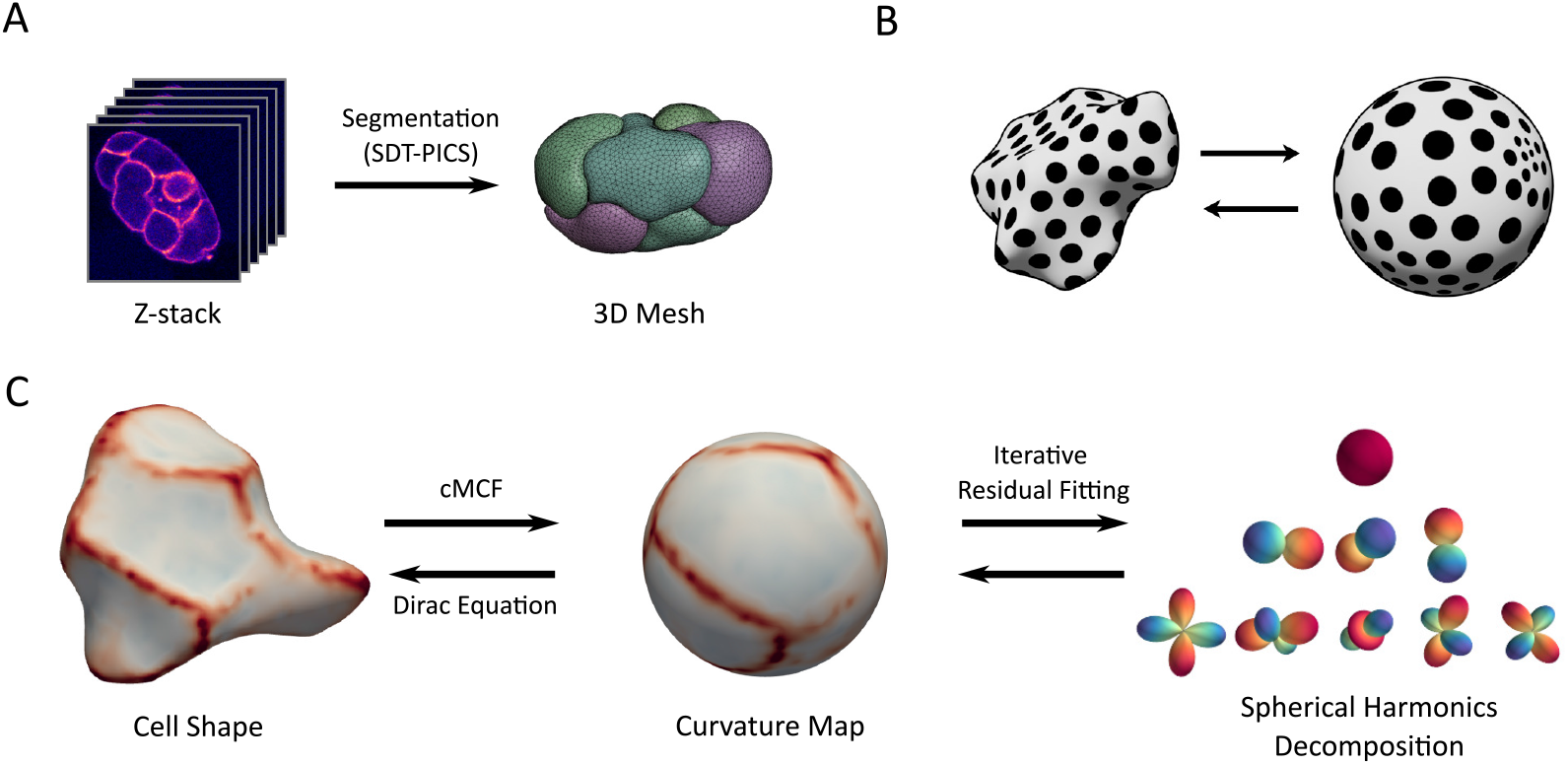
Overview of the ShapeFlow pipeline. A: SDT-PICS segmentation of confocal microscopy timelapses of GFP-tagged embryos results in 3D meshes of the cells. B: Illustration of a conformal map of a shape to the sphere. The map only expands or shrinks the surface without deforming angles. Circles remain circles, and the size of the circles on the sphere represents the local area distortion. C: Overview of the proposed pipeline. In the first step, the shape is mapped to a sphere via conformalized mean curvature flow (cMCF), and can be reconstructed from its curvature function via the Dirac equation. In the second step, the curvature function is decomposed into SH using iterated residual fitting.

A frequency-space representation of shape allows for several efficient analyses. [23] First, SH are intrinsically related to rotations in three dimensions. This allows for an effective and fast alignment algorithm using the fast Fourier transform. Second, because the transform is linear, averaging coefficients also averages the function, which makes it straightforward to calculate an average shape. Third, differences between shapes can be evaluated efficiently, which facilitates distinguishing cells and highlighting shape changes resulting from perturbations. Finally, the SH representation allows for the efficient application of filters on the function, such as smoothing, and the detection of particular shape features.

An overview of FlowShape is given in fig. 1C. As illustrated, the software package also permits reconstructing the shape from the SH representation. To accomplish this, the curvature function is reconstructed from the SH on the sphere, after which the quaternionic Dirac equation is solved. [24–26] Below we go through the steps in more detail. A detailed technical description is included in S1 Appendix.

### Data collection

Live *C. elegans* embryos were harvested from strains which express a membrane-tagged fluorescent protein, either mCherry or eGFP fused to the PH domain of rat PLC1δ (strains OD70 and LP306). They were imaged with laser scanning fluorescent confocal microscopy using a Zeiss LSM 880. The 4D image stacks were segmented with SDT-PICS [20], which outputs a closed triangle mesh representing the cell membranes at each time point.

### Spherical parameterization by conformal mapping

The objective of spherical parameterization is to find a global mapping from the surface to the unit sphere. Many of the algorithms for this approach come from the computer graphics community because such parameterizations are useful e.g. for texture mapping. On triangle meshes, the parameterization gives us a map from the vertices to the sphere by assigning spherical coordinates (*θ, φ*) to each vertex. This mapping necessarily induces distortion: angles, distances and areas as measured on the original surface will not be the same when measured on the sphere. Mapping algorithms can aim to keep the areas or the angles, or both, as close as possible to the original. Here, we use conformalized mean curvature flow [21] to efficiently calculate the spherical parameterization. This flow yields a map that preserves angles, a so-called *conformal* map. The uniformization theorem guarantees that a conformal map always exists, which also holds in the discrete setting. [27]

Once a map is found, the original shape’s surface is represented on the sphere by one or more functions. Previous approaches did this by encoding the x, y and z coordinates of the vertices as three functions on the sphere. [28–30] To simplify our analyses later on, we will use only one function here. It has been shown that given a *conformal* map, the original shape can be fully described using the mean curvature. [31] The function *ρ*, which we will just call the “curvature function,” combines information about both mean curvature and area distortion, expressed here as change in the length of local features, when mapping the surface to a sphere

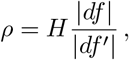

where *H* is the mean curvature. The ratio 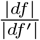 is the local ratio between lengths on the surface and its image on the sphere. Squaring it gives the area distortion.

One issue with conformal maps is that they are not unique. There are many conformal maps from the unit sphere to itself, known as the Möbius transformations. General Möbius transformations on the sphere can be found as compositions of inversions and rotations. Inversions can be understood as fixing the poles that lie on some axis through the origin and then ‘pushing’ the geometry towards one of the poles. To obtain a unique, canonical mapping, we use the Möbius balancing algorithm described by Baden et al. [32]. This algorithm finds the inversion that optimally distributes the area distortion over the sphere. This is beneficial as it facilitates using a minimal number of SH. Further, when trying to find a correspondence between two shapes, we only have to search for the optimal rotation. This is an easier problem than also having to consider the best Möbius transformation to match the shapes.

### Spherical harmonics decomposition

The real SH are a set of polynomial functions on the sphere. Here, we will write the harmonic of degree *ℓ* and order *m* as 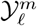. They form an orthonormal basis, which implies that any function on the sphere can be uniquely decomposed as a linear combination of SH.

A real function can be approximated by real SH that are also parameterized by real coefficients. Given a function *ρ* in spherical coordinates (*θ, φ*) the SH decomposition can be written as

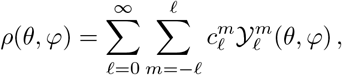

where 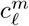 are the real coefficients that need to be found to describe a function on the sphere. While this is an infinite sum, in practice it has to be truncated to a maximum degree *ℓ*_*max*_. This representation is frequency-based and hierarchical. Low degrees capture coarse information and higher degrees capture finer details. The more harmonics are used, the more details are included. For a maximum degree *ℓ*_*max*_ there are (*ℓ*_*max*_ + 1)^2^ coefficients.

To obtain the SH decomposition of a shape’s curvature function, we use the generalized iterative residual fitting (IRF) procedure. This method was originally proposed by Chung et al. [33] and later generalized by Elahi et al. [22]. Here the problem of finding the SH coefficients is efficiently solved by partitioning it into subspaces, one per degree *ℓ*. A least squares problem is solved for each subspace in order, starting with the lowest degree. The algorithm is repeated until convergence.

### Alignment

To compare and average shapes, we want to find a point-to-point correspondence between them. This is known as registration or alignment. Automating the alignment of biological shape data is a non-trivial task, and is an actively researched topic. [34] We give an efficient algorithm to align shapes, by comparing their spherical parameterizations.

The well-known phase correlation algorithm can estimate the translation needed to match two images by finding the maximum of the cross-correlogram. [35] This can be solved efficiently in the frequency domain with the fast Fourier transform. Similarly, for two spherical functions, we can estimate the optimal rotation to align them. Here, the frequency domain consists of the SH.

To align two shapes, they are first mapped to the sphere. We then try to maximize the cross-correlation of their curvature functions *f, g*

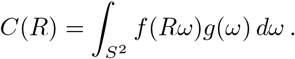

The cross-correlation function *C* is parameterized by a rotation *R*. It can be efficiently computed through a generalized Fourier transform for spherical functions that operates on the SH. [32, 36–38] To evaluate rotations in the space of SH, rotation matrices are calculated using the method described by Pinchon and Hoggan. [39, 40]

This alignment procedure only works pairwise. To align multiple shapes, we propose a simple algorithm that avoids having to calculate all pairwise cross-correlations. We first select one of the shapes as the target, and then align all other shapes to this target. Then, a new target is constructed by averaging all aligned curvature functions. The procedure is then repeated until convergence. The result of applying this procedure can depend on the choice of the initial target, but in practice this did not present any problems for our dataset. To reduce susceptibility to noise, we lightly filter the curvature functions with a small Gaussian kernel before aligning them.

### Shape reconstruction with the Dirac equation

In our framework the inverse problem, i.e. obtaining the shape given the curvature function *ρ* on the sphere, is solved using the quaternionic Dirac equation. [24–26] Locally, a conformal map only rotates and uniformly scales the surface. Quaternions of unit norm are extensively applied to represent rotations in three dimensions. When we allow the norm to be different from one, they additionally define a scaling. This means that we can represent a conformal transformation as a function *λ* : ℳ → ℍ on the surface, taking values in the quaternions.

Below we follow the computational methods developed by Crane et al. [25] and Ye et al. [26]. The quaternionic function *λ* can be obtained from the mean curvature function *ρ* by solving the Dirac equation

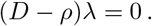

*D* is the intrinsic Dirac operator [26], a differential operator similar to the gradient. In some cases, *ρ* might not correspond to a valid transformation at all. For example, setting *ρ* = 0 everywhere on the sphere is clearly invalid, since it is impossible to remove all the curvature from a sphere. So the equation is relaxed to an eigenvalue problem [25]

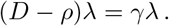

This is then solved for the smallest eigenvalue *γ*.

In the discrete setting, the quaternions act on the edge vectors *e*_*ij*_ of the mesh. Once *λ* is found, new edge vectors are obtained by rotating and scaling them, as prescribed by the quaternions. Obtaining vertex positions from edges then becomes a system of equations

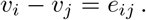

While the Dirac equation guarantees that the system is integrable, numerical errors can still be introduced. Therefore, the system might not have an exact solution. To account for this, the system is solved by least squares.

A more accurate solution can be obtained by iterating the algorithm by varying 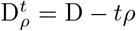 with *t* ∈ [0, 1]. For each iteration, we solve the eigenvalue problem, obtain *λ* and find the new vertex positions with least squares. We increase *t* in equal steps to 1. An accurate solution can be found with three iterations.

Note that this procedure can also be used to flow the surface to a sphere, by setting the curvature to be constant everywhere. However, this is much slower than conformalized mean curvature flow, so we do not use it for this purpose.

To reconstruct a surface, we use a subdivided icosahedron as a starting mesh. This provides a highly regular and uniform sampling of the sphere.

### Implementation and availability

All algorithms were implemented in Python with use of the NumPy [41], SciPy [42] and libigl [43] libraries. Routines for calculating the spherical Fourier transform are provided by the package `lie_learn` by Cohen et al. [44].

## Results

To evaluate our pipeline we used a dataset of around 2,000 cell shapes from 24 *C. elegans* embryo time-lapses. During the nematode’s fast and invariant embryogenesis, cells rapidly change shape, migrate and interact via both signaling and highly specific cellular adhesions. We first looked at limitations and errors induced by our pipeline. Afterwards we evaluated the cells at the seven-cell stage in the wild-type embryo. We compared the cells to each other and evaluated a protrusion filter. Finally, the pipeline was used to evaluate the effects of a gene knockdown on cell shape.

### Validation

All cell meshes were analyzed through our pipeline, expressed in SH, and afterwards reconstructed. We then measured the root-mean-square error (RMSE) for vertex positions of the shapes. We found that the error was 0.26 μm on average, which is low and comparable to the resolution of the microscopy images (± 0.2 μm in the xy-plane). Though the error was low overall, the error was not homogeneous over the shapes. We therefore investigated which features induced higher errors in our pipeline, and how to alleviate the effects. Three sources of error were identified and are discussed in more detail below: (1) discretization effects in areas of high curvature, (2) large area distortion for long thin structures and (3) ridges and sharp features.

We observed that reconstruction errors mostly appeared in areas of high curvature. The discretization of a surface with high curvature results in sharp discontinuities. Because of this, we first investigated the effect of discretization on the quality of reconstruction. We used the Loop subdivision scheme [45] on the meshes. This algorithm recursively subdivides the triangles of the mesh and moves the original vertices based on a spline representation. Each subdivision quadruples the number of vertices. In the limit, this scheme converges to a continuous surface. As we subdivide, the error goes down markedly as shown in fig. 2A. Further, we use the quasi-conformal error 𝒬 [46] to quantify how close to conformal the mapping is. For the spherical mapping, we observe that these 𝒬 -values show linear convergence when the mesh is refined (S4 Fig). However, the reconstruction step is more error-prone. Although the 𝒬 -values improve when the mesh is refined, they plateau at a certain point, indicating a limitation not related to discretization.

**Fig 2.**
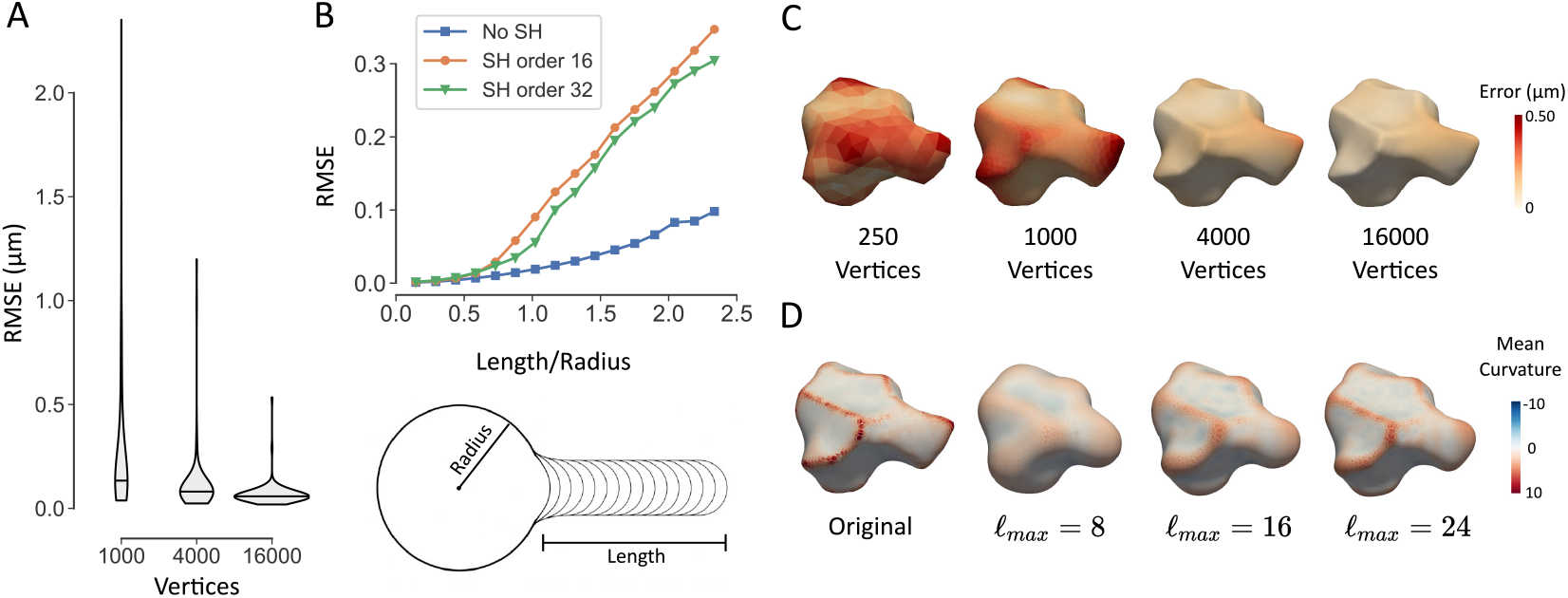
Validation of shape reconstruction. A: Violin plot of root-mean-square reconstruction error (RMSE) of 113 cell meshes. The meshes with 4,000 and 16,000 vertices were obtained by Loop subdivision. B: RMSE for synthetic meshes with a single protrusion of variable length. The length of the protrusion is expressed relative to the radius of the sphere. The RMSE is shown for three cases: reconstruction with SH decomposition for a maximum order of 16 and 32, and reconstruction without SH decomposition. C: Reconstruction error shown on one of the wild-type ABpl cell meshes at various resolutions. D: Influence of maximum SH order on the quality of reconstruction for an example ABpl cell mesh.

Because the spherical parameterization is conformal, the area distortion can locally be very large. This is rare in our data, but it can be a problem for shapes with long thin structures, such as neuronal axons. Long protrusion-like structures will be mapped to a small part of the sphere, where the curvature function would vary considerably over short distances in these regions. To test the algorithm in this situation, a synthetic dataset was made, consisting of a sphere with a single protrusion of variable length (fig. 2B). When the protrusion becomes longer than the radius of the sphere, the reconstruction error starts to become larger (fig. 2C). This effect is much less pronounced when the SH decomposition is skipped. Hence, it is mostly the SH decomposition that fails to capture the detail in this small region. Without SH decomposition, some meshes may still fail to be accurately reconstructed, as illustrated on synthetic examples shown in supplementary S1 Fig. In these cases, the area distortion (measured as the ratio of triangle areas) on the spherical map was severe, up to 10^−8^.

Figure 2D shows the effect of the maximum degree *ℓ*_*max*_ of SH. A lower maximum degree needs fewer coefficients to represent the shape, at the cost of a loss in high frequency details, such as ridges and sharp features. The average power spectrum (see S1 Appendix section 3.4) shows that there is increasingly less information in the higher degrees (S3 Fig). Given that each degree *ℓ* introduces an additional 2*ℓ* + 1 coefficients, it is clear that after a certain point there will be diminishing returns. For our analyses below, we used a maximum degree of 24, which captures enough detail to faithfully reconstruct the shape (see also fig. 2D).

### Wild-type embryo

We next applied the pipeline to further analyze our dataset of cells in the early *C. elegans* embryo. As the development of *C. elegans* is invariant, each cell in the lineage has a unique name. We focus here on the seven-cell stage, which occurs directly after the division of the EMS cell into E and MS. The other cells at this stage are ABal, ABar, ABpl, ABpr and P2. First, we wanted to quantify cell shape changes over time. The sum of squares of the SH coefficients results in the total Willmore energy 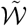, a measure of how much the cell deviates from a sphere. We followed changes in 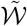 over a cell’s lifetime (S5 Fig). Overall, cells start out relatively spherical (just after division), obtain a maximum energy in the middle of their life and then become spherical again before dividing. There is an uptick in 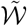 at the final step, reflecting the appearance of the cleavage furrow. Individual cells also show markedly different behavior, with ABpl and MS having relatively high values, marking active behavior and cellular movement, whereas P2 and ABal retain more spherical shapes.

We next evaluated whether cells could be distinguished based on shape alone. To do this, we took the shape averages, by averaging the SH coefficients, over time for all individual cells of the seven-cell stage. As we infer from the results above that cells can be most easily distinguished in the middle portion of their lifetime, we did not use the first and last quarter of a cell’s lifetime when calculating the average shape. We further assume that cells do not rotate much during their lifetime, so no alignment was applied.

Next, the normalized correlation coefficient for every pair of shapes was calculated using our alignment procedure (see below). This correlation coefficient was then transformed into a distance measure by 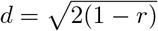. We then applied the UMAP dimension reduction algorithm [47] on the distance matrix obtained to get a two-dimensional latent representation (fig. 3A). For three clusters, average cell shapes are shown, illustrating the shape changes that distinguish them.

**Fig 3.**
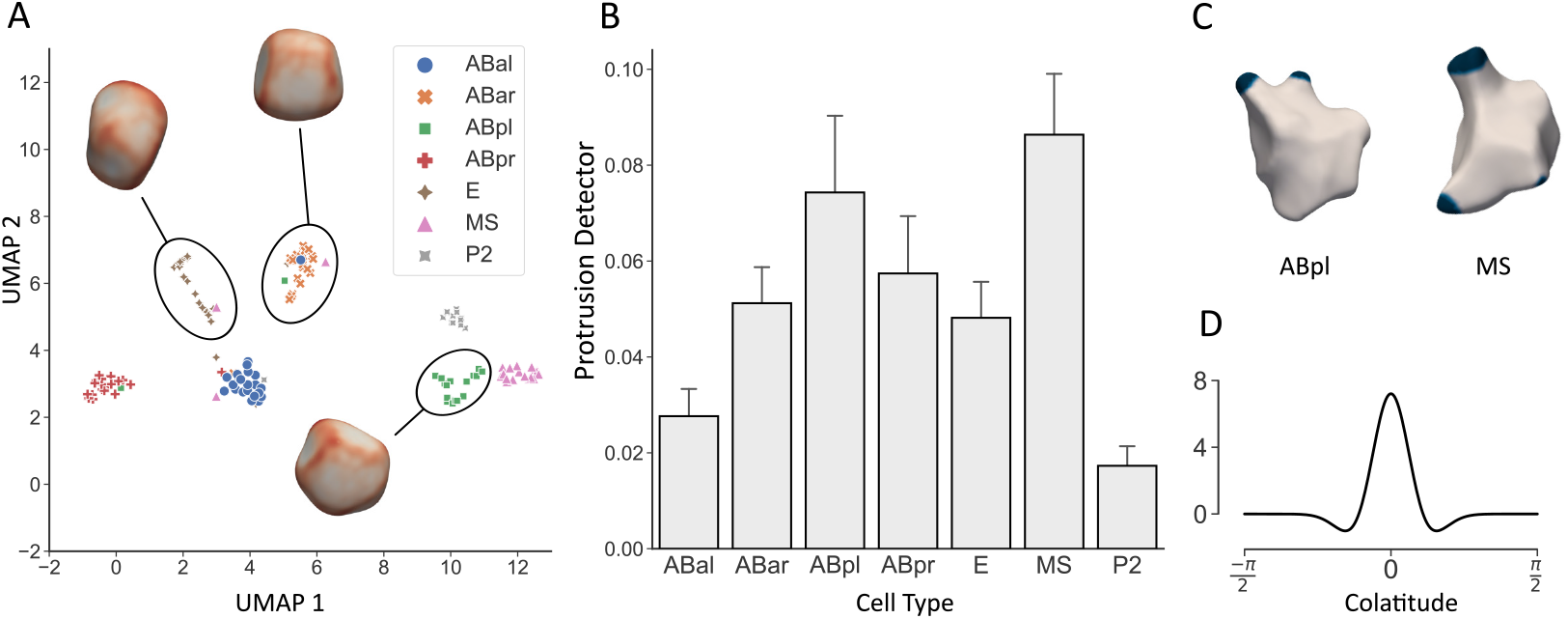
Results for the wild-type seven-cell stage embryos. A: Result of UMAP dimension reduction of cell shapes from the seven-cell stage. Some selected average shapes are shown for clusters detected with *k*-means clustering. B: Results of the protrusion detector for seven-cell stage. C: Two example cells with detected protrusions. Blue color indicates where the Laplacian of Gaussian filter is above threshold. D: The filter kernel plotted as a function of colatitude *θ* (angle from the north pole of the sphere).

To check the accuracy, we ran a *k*-nearest neighbor classification (*k* = 5) on the low dimensional representation with 10-fold cross-validation, resulting in a mean accuracy of 91.3%. Note that this clustering is based only on shape, and is invariant with respect to position, rotation and scale of individual cells. We conclude that at least for early embryogenesis, cell shapes are highly consistent, and cells can be accurately distinguished by their shape alone.

One important active behavior of cells in the early embryo is the development of protrusions such as lamellipodia. One application of the presented framework is to automatically detect such features using a custom filter. This is analogous to ‘blob detection’ in image analysis. [48] We use a Laplacian of Gaussian filter (fig. 3D) to detect blobs in the curvature function, and then use a simple threshold. The total area above threshold is then used as a proxy for the existence of protrusions such as lamellipodia. Figure 3B shows the result of this filter to all shapes from the seven-cell stage. ABpl and MS score highest, but other cells also often show protrusions, which confirms previous findings. [49] ABpl indeed develops striking lamellipodia (fig. 3C), and both cells actively change shape and move during their lifetime. [49]

### Phenotyping of genetic perturbation

Wnt signaling plays a central role in determining cell fate in the early embryo, while it also has important roles later in development, such as in neuronal cell migration. [50] For the endomesodermal precursor cell (EMS), a Wnt signal polarizes division at the end of the four-cell stage. The asymmetric division that follows results in differentiation of the endodermal lineage with the birth of the endodermal founder cell E. It also induces a lasting rearrangement of the cortical actomyosin in this cell. [4] Removing DSH-2 and MIG-5 by RNAi knockdown prevents this rearrangement. DSH-2 and MIG-5 are both Dishevelled (DSH) proteins, a family of proteins that is involved in both canonical and non-canonical Wnt signaling. Here we use our pipeline to determine whether there is a significant difference in the shape between cells in wild-type and dishevelled RNAi embryos.

To compare the shapes between the conditions we first calculate the average cell shape per condition. First the cells are aligned, after which the average of their curvature functions is taken.

To then compare the average shapes 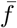 and 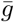, we reconstruct the shapes and next use as a distance measure the root-mean-squared distance between the surfaces

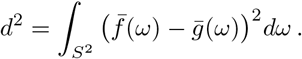

To calculate the empirical distribution 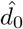 under the null hypothesis that there is no difference between average shapes, we randomly assign the shapes to each group and calculate *d* as above, repeating this 1,000 times. We can then estimate 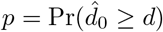. Rewriting this gives 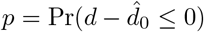, shown on fig. 4A. To adjust for multiple testing, we apply a Bonferroni correction.

**Fig 4.**
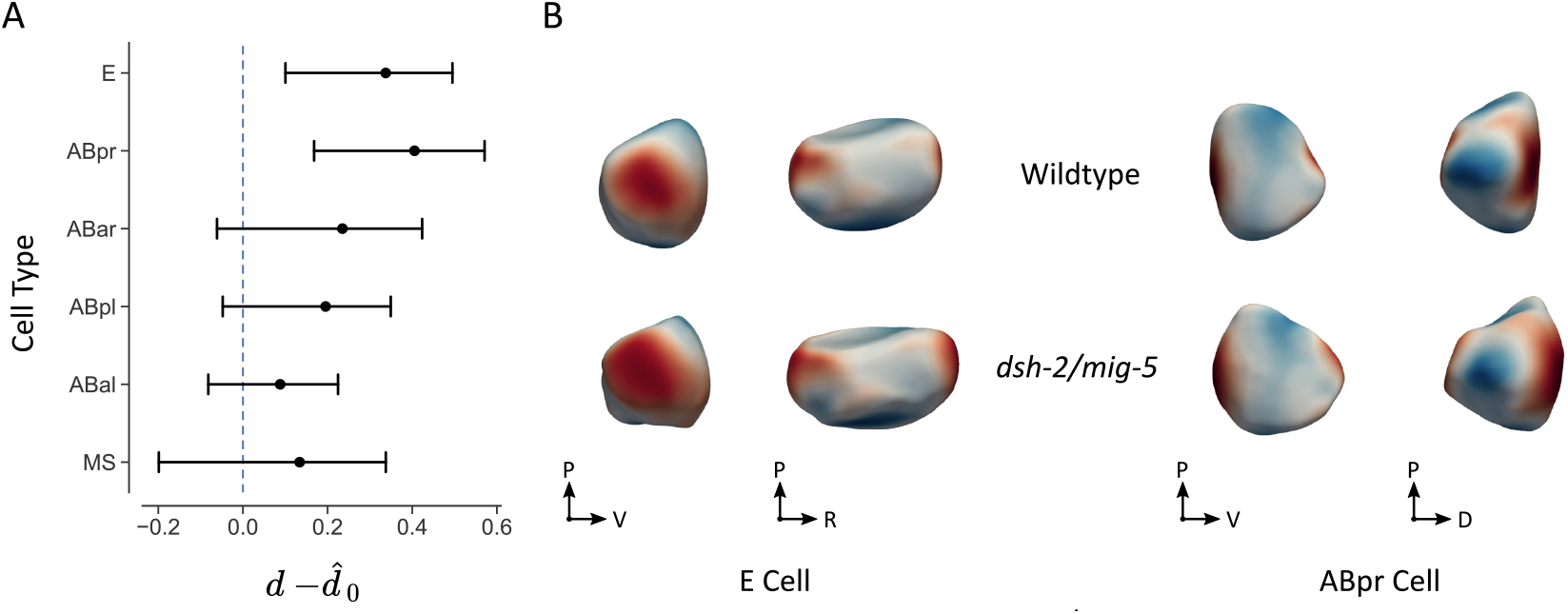
Differences found between wild-type and *dsh-2/mig-5* RNAi knockdowns. A: 95% confidence intervals for difference 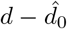 between the average shapes of wild-type versus *dsh-2/mig-5* knockdowns. After Bonferroni correction, we find a significant difference for E (*p* = 0.024) and ABpr (*p* = 0.020). The P cell is left out of the analysis as it undergoes division during this stage. B: Reconstructed shapes for the averages of E and ABpr cells. Color signifies difference along the surface normal between wild-type and perturbation, with red indicating outward displacement in the perturbed cells and blue indicating an inward displacement.

We find a significant difference for E (*p* = 0.024) and ABpr cells (*p* = 0.020). The observed shape changes are plotted in color code on the average shape (fig. 4B). We conclude that E cells in the *dsh-2/mig-5* RNAi knockdown become more elongated and indented, together with the observed altered cortical actomyosin. Also, the ABpr cell appears to be more compressed by its neighbors. This cell lies next to E, which may explain the shape change; no direct link to Wnt signaling is currently documented.

### Performance

For a mesh of 1,000 vertices, the whole pipeline takes about 2 seconds. All tests were done on a Windows laptop with an AMD Ryzen 3550H processor at 2.1 GHz. Table 1 and S2 Fig show a breakdown of the time taken by different steps. Spherical parameterization and SH decomposition scale roughly linearly with mesh density, while the reconstruction step scales quadratically. For larger meshes the reconstruction step quickly dominates. Note that during analysis, the reconstruction step is not of importance, so it can be skipped. The spherical Fourier alignment procedure takes 70 ms on average, and evaluating the rotation matrices for the SH around 5 ms (*ℓ*_*max*_ = 32 for both).

**Table 1.** Time (in seconds) taken by the different steps in the pipeline. SH decomposition uses *ℓ*_*max*_ = 32. See S2 Fig for more detail.

| Vertices | 1,000 | 5,000 | 10,000 |
| --- | --- | --- | --- |
| Spherical parameterization | 0.37 | 1.5 | 4.0 |
| SH decomposition | 1.1 | 4.8 | 9.0 |
| Reconstruction | 1.7 | 27 | 132 |

## Discussion

We have presented FlowShape, a framework that can be applied to analyze cell shapes. First, the mean curvature of the shape is conformally mapped onto a sphere. The procedure to generate this map is parameter-free and does not require any landmarks. The curvature function is unique and can completely represent a shape in a parsimonious way. This function is then decomposed into a weighted sum of SH. We have shown that this representation allows for aligning, averaging and comparing shapes, and enabled us to effectively distinguish and characterize early embryo cells. Further, we used the Willmore energy to characterize cell shape change over time and designed a filter to highlight narrow extrusions from the cell, such as lamellipodia. Finally, we were able to leverage the representation to compare perturbed to wild-type cells in a sensitive manner, and illustrate the changes in shape.

By only using the mean curvature to represent shape, shape alignment can be done via the fast Fourier transform procedure. This procedure is highly efficient, does not require an initial guess and always finds the globally optimal solution for rotation alignment of the shapes. Previous approaches encoded the shapes on the sphere using three coordinate functions [28–30], which can complicate some analyses. These coordinate functions are not independent, and they “mix” when rotated, which makes aligning the maps of different shapes more complicated. To align shapes using this representation, the contribution of the SH of degree *ℓ* = 1 has been used, as this is the first order ellipsoid (FOE). [51] The ellipsoid is then used to define a canonical orientation, as it has three perpendicular axes. We identify two shortcomings of this method. First, an ellipsoid does not actually define a unique rotation, as it is symmetric to 180° rotations around each of its axes. Second, when two of the major axes are of similar size, as is often the case, the found orientation will be sensitive to noise. As an illustration, S6 Fig shows a comparison between the proposed cross-correlation and FOE alignment where the latter fails to properly align features.

Our alignment approach also contrasts with a commonly used algorithm to align shapes, the Iterative Closest Point (ICP) algorithm [52], as this procedure only guarantees a locally optimal solution.

Our method is computationally very efficient, as we only solve linear systems, most of which are sparse. The performance of comparable approaches as reported in the literature is many times slower, as seen, for example, in the article by Shen et al. [11], where the nonlinear optimization method of Brechbühler et al. [51] was used to calculate spherical mappings. They report that spherical parameterization per mesh with around 2,500 vertices typically took 15 minutes to 3 hours.

The SH description of shapes given by FlowShape involves numerous descriptors, as shapes are represented completely and accurately. This is a trade-off to using simple shape descriptors, such as sphericity, that can capture particular shape features. Further, the SH functions by themselves are difficult to interpret, as they are global functions, and work together to represent local features. One way to deal with this issue is to apply filters such as edge detectors to mark local features. Here we designed a protrusion detector to illustrate the idea. Applying filters can be done very efficiently in the SH representation. One limitation in the current setup is that this filter has to be axially symmetric, which limits the features that can be searched for. It is future work to expand our toolbox to work with any possible filter. Further, as the frequency-space features obtained from the spherical harmonics decomposition allow for efficient filtering, they can be used directly in machine learning methods such as convolutional neural networks (CNNs). [38] Our approach can therefore readily be used in fields where such methods are applied, for example to classify cells by their shape.

We showed that the pipeline achieved high accuracy on our dataset. Though the reconstruction algorithm works well in practice, it may perform worse for some shapes, especially those with long thin structures. Such structures are mapped to very small areas on the sphere and can not be accurately reconstructed with the method we used here. We can improve reconstruction quality to a point by increasing mesh density, at the cost of computational time. To further mitigate the issue, an area correction term can be added to the reconstruction, as shown by Ye et al. [53]. As high area distortion was not a significant issue for our dataset, we did not pursue this approach here for simplicity as it would imply tracking two functions instead of just one. It should be noted that reconstruction is not required for most SH-based analyses of cell shapes.

In this paper we successfully used average shapes in order to, for example, compare perturbed cells to wild-type. However, averages may be too simplistic in some cases as they do not capture the variable parts of shapes in a population of cells. For example, a sharp ridge that slightly varies in position between observations would be represented as a broad, blunt ridge on the average shape. A more expressive representation may be needed to make a generative statistical model in such cases. This is not a trivial exercise, as adding variance is not a linear operation and variance of the curvature functions is not equivalent to the variance of the SH coefficients. Random fields present a promising approach to solving this problem [54], which is left for future work.

Cell shapes are tightly controlled, a hallmark of polarization, differentiation and cellular activity. Linking shapes to the processes that control them is an area of active research. To facilitate this research, we propose FlowShape as an efficient and powerful tool to fully characterize, compare and distinguish cell shapes. The pipeline can play a role in recovering more quantitative information from microscopy images, or applying machine learning to the analysis of microscopy data. Further, the pipeline is generic in design, and can therefore be applied more broadly. In essence, any shape that has a closed surface with no holes (genus 0) can be analyzed. Hence, apart from cells, FlowShape can be applied to the analysis of other shapes in the life sciences, including tissues or structures at the anatomical level.

## Supporting information

**S1 Appendix. Technical details**. A more detailed look at the mathematics behind FlowShape. It includes implementation details as well as some mathematical derivations.

## Acknowledgments

Experiment expenses were supported by FWO research grant G055017N, W. Thiels was supported by FWO fundamental research grant 11I2921N and C. van Bavel by FWO fundamental research grant 11L0923N.

## Supplementary figures

**Fig S1.**
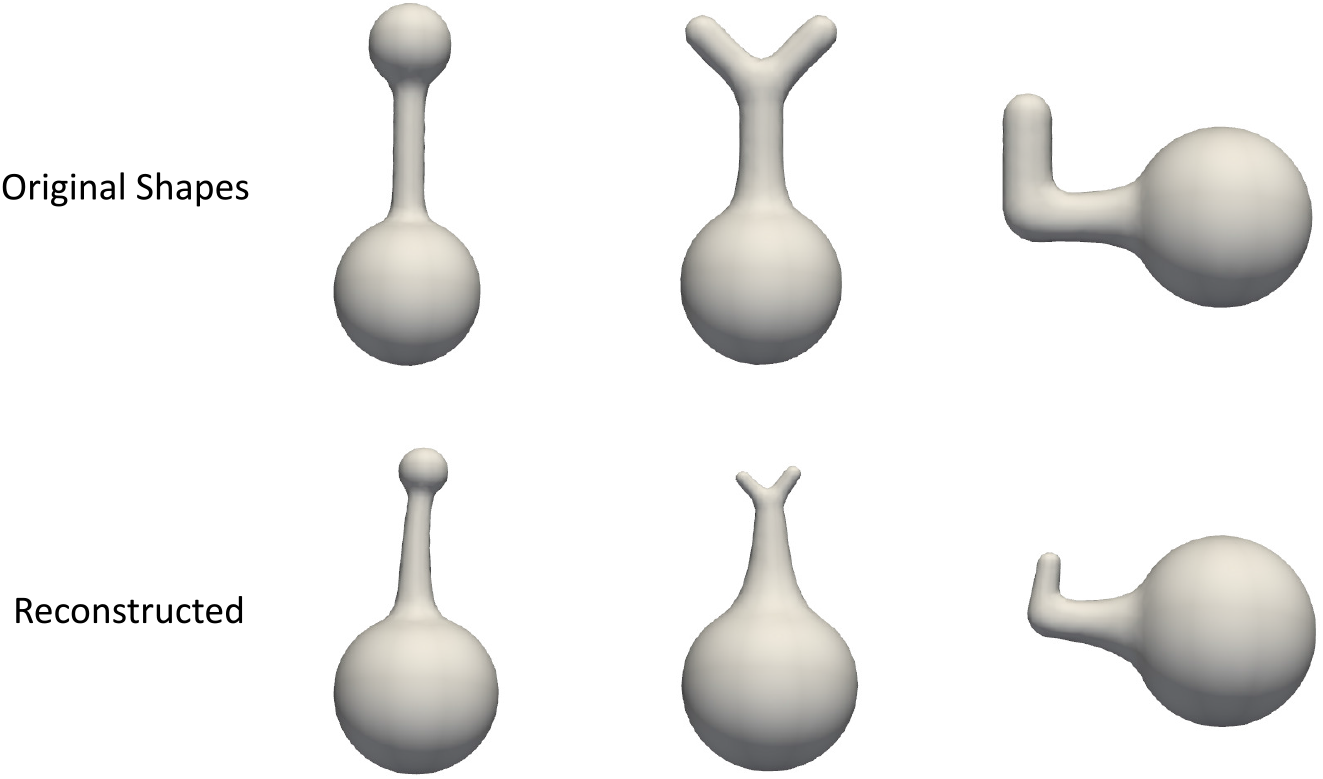
Synthetic shapes with high error. These reconstructions were made directly from the conformal mean curvature map on the sphere, without SH decomposition.

**Fig S2.**
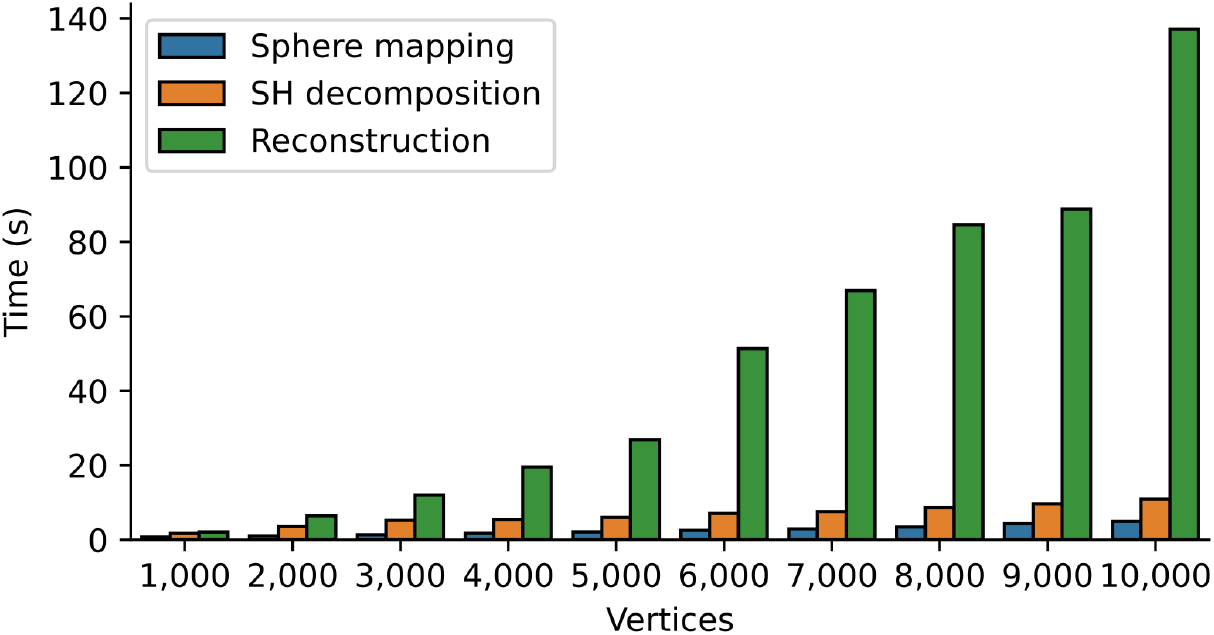
Performance versus mesh density. Wall-clock time in seconds for the different steps in the pipeline for meshes of varying sizes. SH decomposition uses *ℓ*_*max*_ = 32. The reconstruction step quickly dominates, but it is not necessary for most of the analysis.

**Fig S3.**
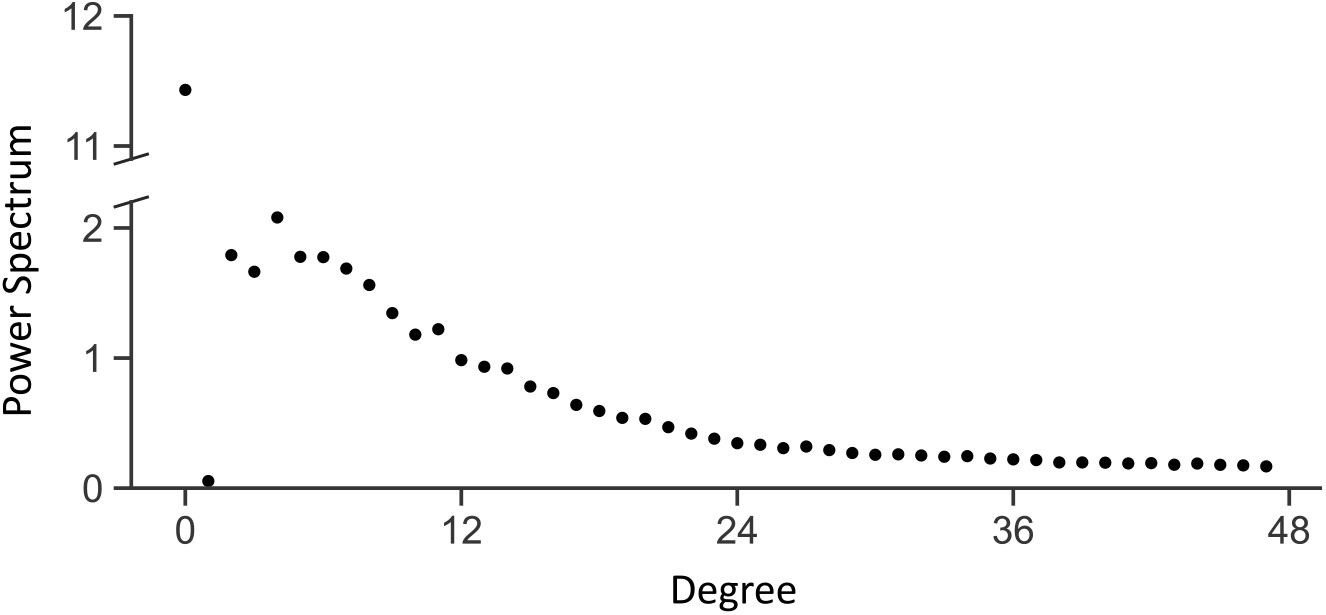
Average power spectrum. Average power spectrum of the curvature function for our dataset of wild-type *C. elegans* embryo cells. The norm of the first degree is very small because it corresponds to Möbius transformations, which are removed by the Möbius balancing.

**Fig S4.**
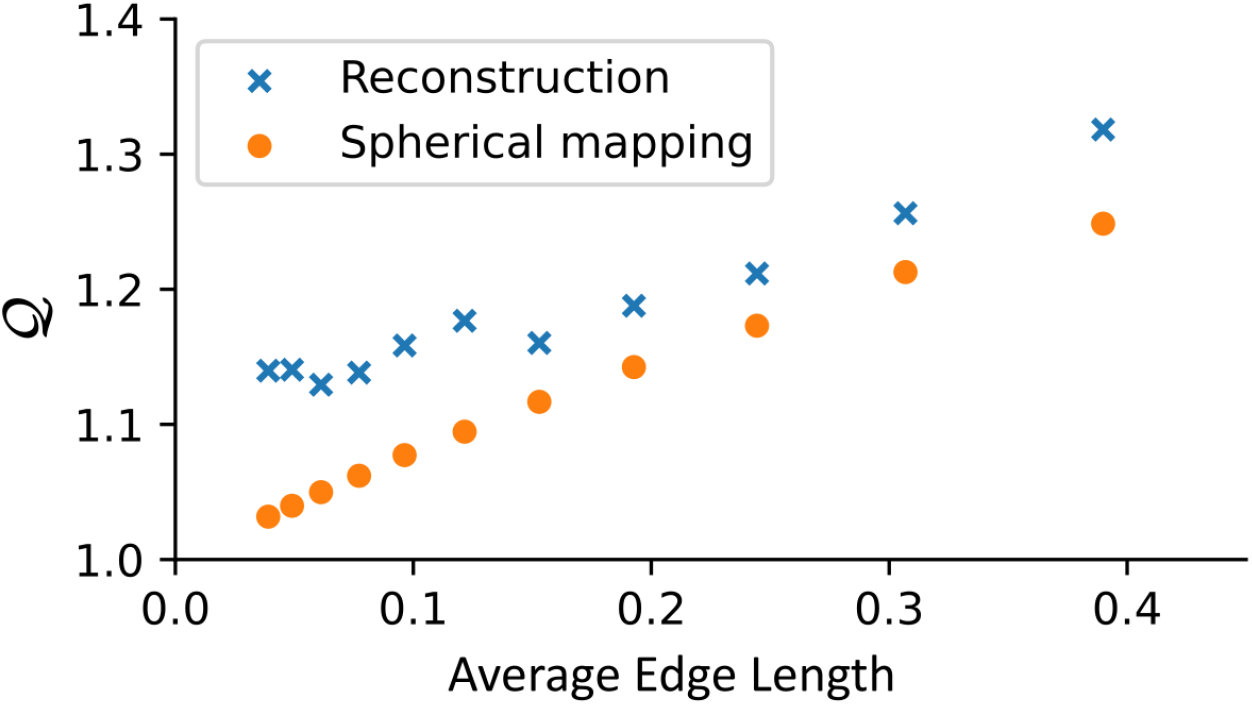
Convergence of 𝒬-values. Average 𝒬-values versus average edge length for various mesh densities. A 𝒬-value of 1.0 represents a perfectly conformal transformation. Higher densities give lower overall error.

**Fig S5.**
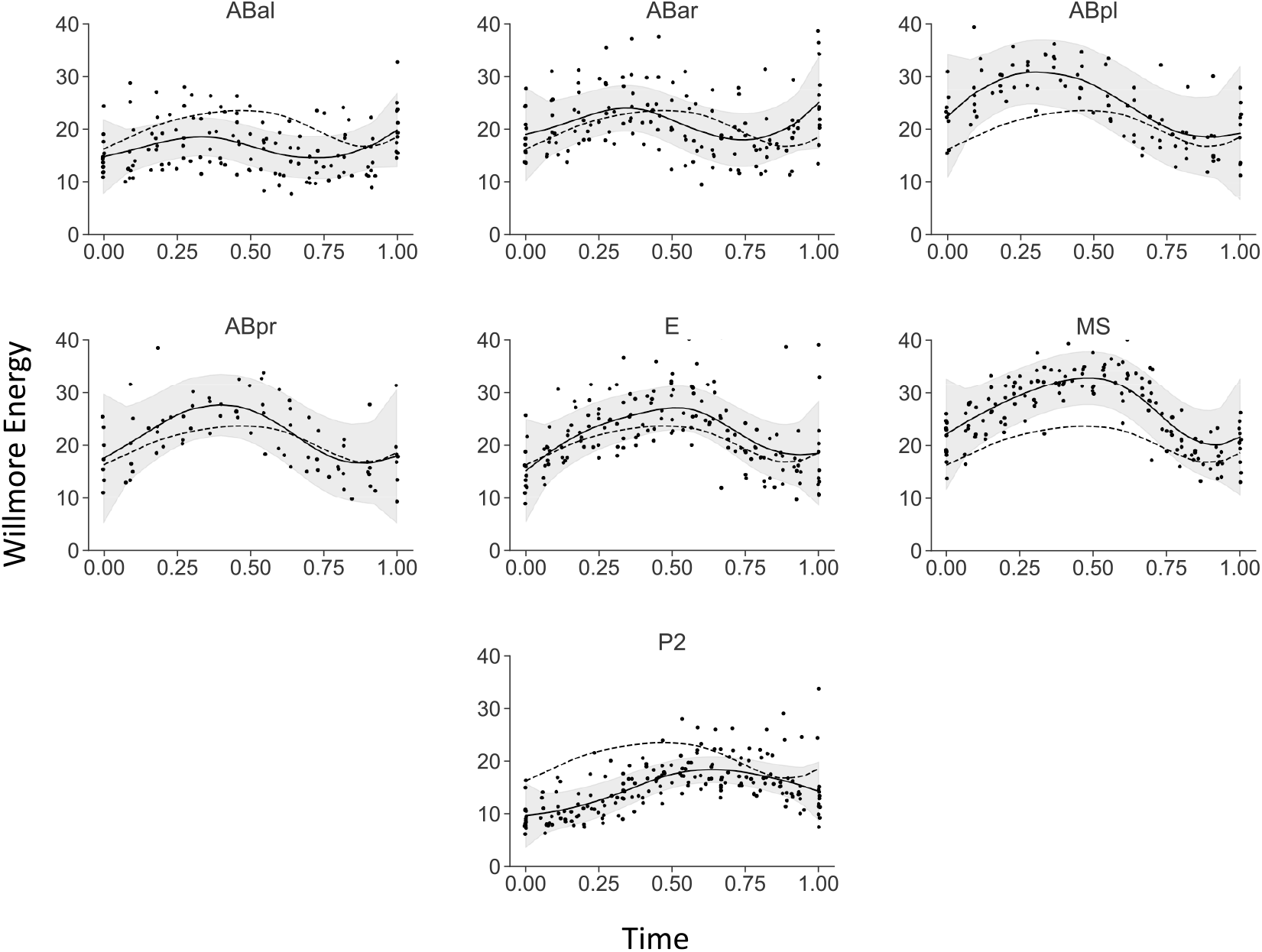
Willmore energy over the lifespan of cells. We remap the time that a cell is ‘alive’ (between divisions) to the interval [0, 1]. Solid lines show LOWESS fit (with 90% CI) of Willmore energy 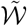 over this lifespan of cells in our *C. elegans* embryo dataset. Dashed line shows the average for all cells.

**Fig S6.**
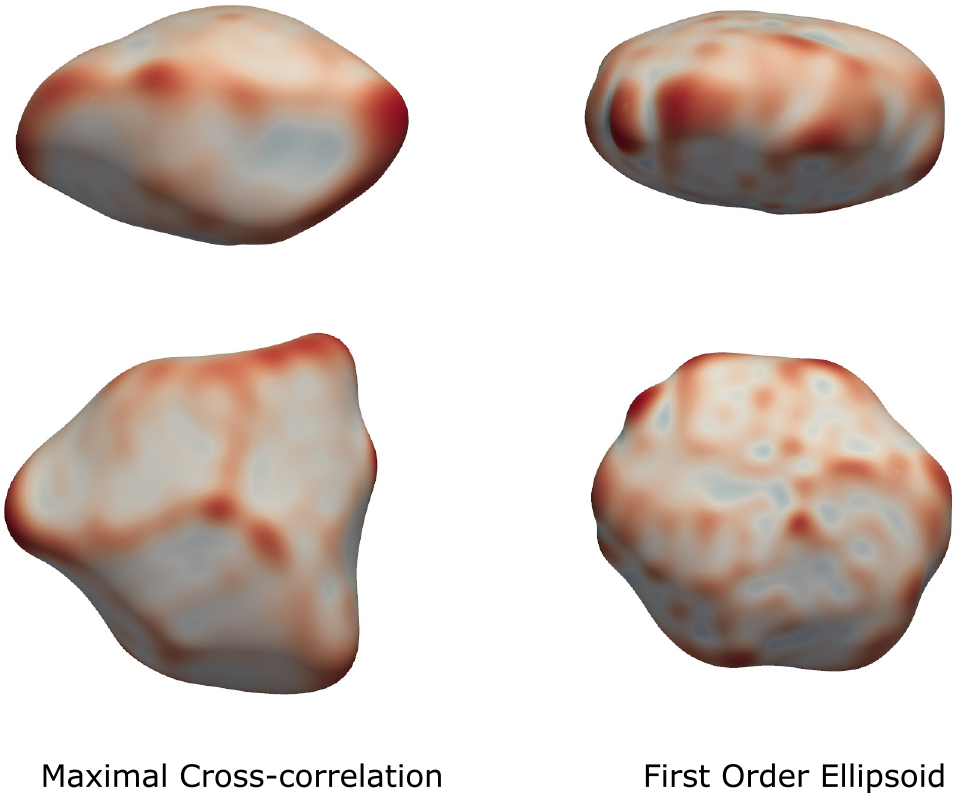
Comparison of the average shapes obtained with our method and the first order ellipsoid. Comparison of the average shape for the ABpl cell. Left: Average after alignment with maximal cross-correlation (our method). Right: Average after alignment with the first order ellipsoid. Averaging the cell based on the ellipsoid loses characteristic features of the cell that are maintained by our alternative approach.

## S1 Appendix: Technical details

### 1 Spherical harmonics

As we represent the shape as a scalar function on the sphere using *ρ*, we can apply concepts from spectral analysis to parameterize the shape. The *spherical harmonics* (SH) decomposition is the analogue of the Fourier transform, applied to a spherical domain. It decomposes a function into an infinite sum of basis functions. This decomposition can be found if *ρ* is square-integrable

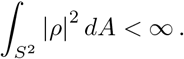

The real SH form an orthonormal basis of the space of square-integrable functions 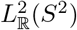. This means that every square-integrable function 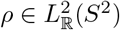 can be decomposed as a linear combination of SH.

Usually, the SH are given as complex functions, but they can be transformed into a real-valued form. For a real function, using the real SH has the added benefit that the coefficients are also guaranteed to be real.

Since *ρ* is a function on the sphere, we can parameterize it with spherical angular coordinates (*θ, φ*), with *θ* the colatitude (0 ≤ *θ* ≤ *π*) and *φ* the longitude (0 ≤ *φ <* 2*π*). The SH decomposition is

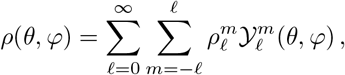

where 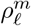 are real coefficients and 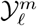 are the SH.

#### 1.1 Definitions

The SH are a set of functions defined on the surface of a sphere. The complex SH 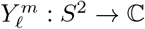 of degree *ℓ* and order *m* are defined as

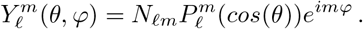

*N*_*ℓm*_ is the normalization factor

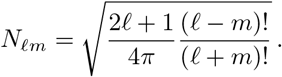

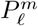 are the associated Legendre polynomials, which can be defined in terms of the ordinary Legendre polynomials *P*_*ℓ*_(*x*)

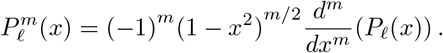

The ordinary Legendre polynomials can be expressed as

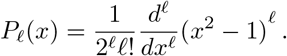

The real SH 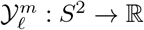 can then be derived as follows

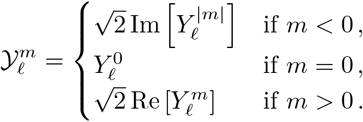

For every degree *ℓ* there are 2*ℓ* + 1 such functions, with order *m* = −*ℓ*, …, +*ℓ*.

The orthonormality of SH means that the inner product satisfies

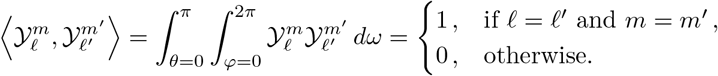

#### 1.2 Implementation

Since *ρ* is a function on the faces, we will take the spherical coordinates of the triangle barycenters to be the sampling points

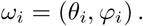

The barycenter of a triangle is calculated by taking the mean of the Cartesian coordinates of the three vertices. Since this mean does not lie on the sphere, we project it back to the surface.

We use the generalized iterative residual fitting (IRF) procedure, originally proposed by Chung et al. [33] and later generalized by Elahi et al. [22]. In this method, the problem of finding the SH coefficient is efficiently solved by partitioning the problem into subspaces. Here, we use one subspace per degree *ℓ*. Let *c*_*l*_ be the column vector of estimates with length 2*ℓ* + 1, and Y_*ℓ*_ a matrix with size *n* × (2*ℓ* + 1)

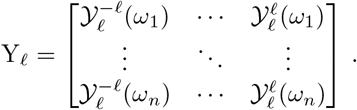

Set the initial residual *r*_0_ = *ρ*. The first subproblem is then to find *ĉ*_0_ that minimizes

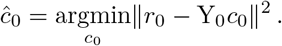

The estimate is Y_0_*ĉ*_0_ so that the new residual is

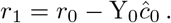

This procedure continues iteratively. At step *j* we find *ĉ*_*j*_ that minimizes

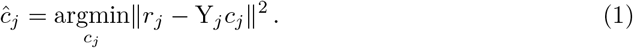

Updating the residual with

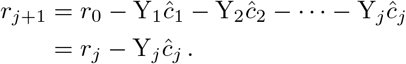

At each iteration *j*, eq. (1) is solved using weighted least squares

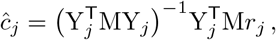

using the diagonal mass matrix M as weights to account for unequal triangle sizes. The estimate for *ĉ* is then just the concatenation of all the *ĉ*_*j*_. This whole procedure is then iterated again, until the norm of the residual can not be decreased further.

### 2 Fourier methods and alignment

The convolution theorem says that we can find the convolution of two functions *g* and *h* by simply multiplying them in the Fourier domain. We will use this principle in two ways: first, to efficiently evaluate filters on spherical functions, and second, to calculate cross-correlations.

#### 2.1 Filters

Given kernel *h* and a function *f*, the convolution is [55]

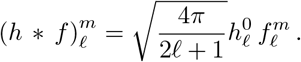

Note that the kernel only has coefficients where *m* = 0, which is equivalent to saying it is axially symmetric. For example, a smoothing filter can be expressed by the heat kernel *G*_*k*_

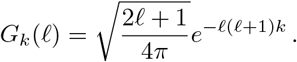

This heat kernel is the spherical analogue of the well-known Gaussian distribution. [56]

The Laplacian of Gaussian is often used to filter for 2D images. [48] It is straightforward to derive an analogous filter for shapes

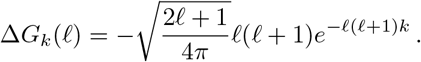

By doing the filtering on the spherical domain instead of the original mesh, the conformal scale factor is ignored. The result is that the filter kernel is locally scaled, but because the map is conformal, it remains isotropic.

#### 2.2 Aligning shapes

The inner product for two functions *f, g* ∈ *L*_2_(*S*^2^) is

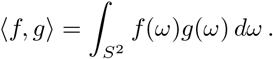

Because of the orthonormality of SH, this can also be expressed as the inner product of their SH coefficients 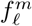 and 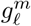

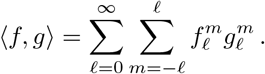

Given two shapes, we want to find the rotation that best aligns them. To do this we map them both to the sphere and calculate their curvature functions *f* and *g*. We then try to maximize their cross-correlation

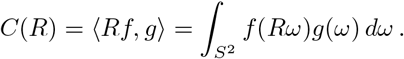

This is similar to a convolution except that the argument *R* is a rotation, so *C* is a function from the space of rotations SO(3) to ℝ. We want to find a rotation *R* that maximizes this cross-correlation. Evaluating *C*(*R*) on all possible rotations would be extremely costly. Fortunately, this can be efficiently computed thanks to a Fourier transform. [32, 36, 37] Here the forward transform is the SH decomposition. The inverse transform is done using the generalized Fourier transform over SO(3). The element in the Fourier domain of SO(3) is formed by taking the outer product of each subspace *ℓ* of the SH, forming a sequence of (2*ℓ* + 1) × (2*ℓ* + 1) matrices. Rotations are parameterized using ZYZ Euler angles (*α, β, γ*), with *α, γ* ∈ [0, 2*π*) and *β* ∈ [0, *π*). A rotation matrix is thus expressed as R = R_*z*_(*γ*)R_*y*_(*β*)R_*z*_(*α*)

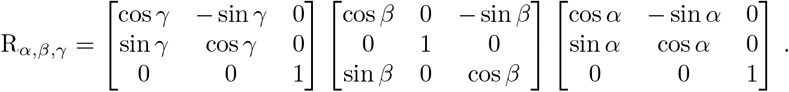

The output of generalized Fourier transform is a 2*B* × 2*B* × 2*B* cube of Euler angles (*α, β, γ*). *B* = *ℓ*_*max*_, the band-limit and each point has a value *C*(*R*_*α,β,γ*_).

After finding the grid point that maximizes the correlation, we do a local quadratic fit to estimate the location of the peak at sub-grid precision. At a bandwidth of *B* = 32, the grid is sufficiently densely sampled so that the average error is about 1°.

To evaluate rotations efficiently in the space of SH, rotations matrices are calculated using the method described by Pinchon and Hoggan [39, 40].

### 3 Mean curvature and the Dirac equation

We assume a shape to be analyzed is a differentiable two-dimensional manifold ℳ. The manifold has an immersion in ℝ^3^

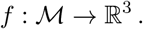

The differential *df* of this map is a linear map, from the tangent space at every point to tangent vectors in ℝ^3^. Analogous to the Jacobian matrix, it gives the best linear approximation to the surface at a point.

We also require that the surface is topologically equivalent to the 2-sphere, so it is simply connected, without boundary and of genus 0.

The Bonnet problem is concerned with the existence of surfaces that have the same mean curvature at corresponding points. In general, there exist so-called Bonnet pairs. These pairs of surfaces are related by an isometry but have the same mean curvature at corresponding points. [57, 58] For surfaces with spherical topology, this is not the case. A proof that these can be uniquely described by their mean curvature is given by Lawson and de Azevedo Tribuzy [31].

#### 3.1 Quaternions

The algorithms discussed here rely on some quaternionic calculus. We refer the reader to Vicci [59] for an overview of quaternions and only review the necessary properties. The quaternions ℍ are an extension of the complex numbers, where three basis quaternions **i, j** and **k** take the place of the imaginary unit *i*. A quaternion can be represented as

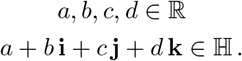

We identify vectors in ℝ^3^ with the *imaginary* quaternions

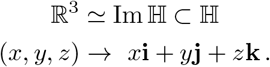

So that we can write any quaternion as a scalar plus vector part: *q* = *a* + **v**. It is well known that quaternions can be used to represent rotations in three dimensions. To be specific, let *q* be a unit quaternion (|*q*| = 1). Then q represents a rotation of the vector *u* as

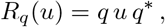

where *q*^*^ denotes the conjugate quaternion. If *q* has a magnitude different from 1, then *q u q*^*^ represents a rotation and uniform scaling. Writing *q* as its scalar and vector parts,

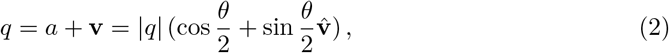

the uniform scaling factor is |*q*|^2^, the rotation angle is *θ* and the rotation axis is the normalized vector 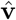.

#### 3.2 Spin transformations

The immersion 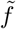 is called a spin transformation of *f* if there exists a smooth quaternion-valued function *λ* : ℳ → ℍ such that

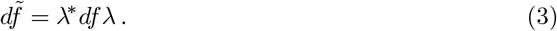

This relation is conformal by construction, as the tangent spaces are transformed by a local scaling and rotation as in eq. (2). The area is locally scaled by

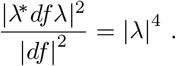

We require that the surface is topologically equivalent to a sphere. This guarantees that any two conformal immersions are spin equivalent. [31, 57] By the uniformization theorem, a conformal map always exists. This also holds in the discrete setting. [27]

#### 3.3 Dirac equation

Not every function *λ* corresponds to a valid spin transformation, as the differential *λ*^*^*df λ* might fail to be integrable. In order for *λ* to be integrable it has to satisfy the Dirac equation

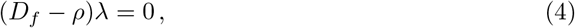

where *ρ* : ℳ → ℝ is a scalar function and *D*_*f*_ is the extrinsic Dirac operator, as originally described by Kamberov et al. [57].

Solving eq. (4) by prescribing some *ρ* gives a new immersion 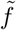. The mean curvature functions *H* are related by

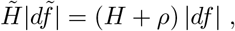

where |*df* | is the length element. The intrinsic Dirac operator *D* is defined as [26]

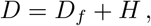

such that solving the intrinsic Dirac equation

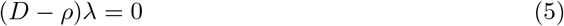

gives a new immersion 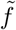 as above. The mean curvature is related by

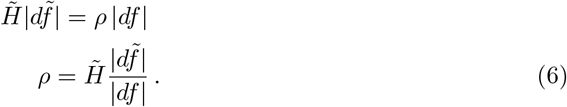

This gives an effective algorithm for recovering a shape from a sphere: save the function *ρ* as in eq. (6) on the sphere, solve the intrinsic Dirac equation eq. (5), and recover 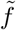 by integrating 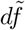. This operation is conformal by construction, so it only works if the original spherical map is conformal.

#### 3.4 Willmore energy and the power spectrum

The Willmore energy of a surface ℳ is the squared norm of the mean curvature [60]

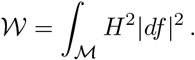

Since we are dealing with closed surfaces of genus 1, we have 𝒲 ≥ 4*π*. When ℳ is exactly a sphere, 𝒲= 4*π*. For this reason, some authors use 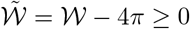 instead. This quantity can be used to measure how much a surface deviates from a perfect sphere. It is also invariant under Möbius transformations. [61] also has a physical interpretation, as for a thin elastic sheet (i.e. the cellular cortex), 𝒲 is proportional to the total bending energy. [62]

If we have the conformal map to the sphere, we want to find an expression for 𝒲 in terms of the curvature function *ρ*. Squaring both side of eq. (6), now with |*ds*| as the length element on the sphere yields

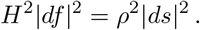

The area scaling cancels out, so that we can evaluate the Willmore energy directly on the sphere

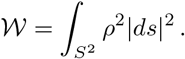

If we have the SH decomposition of *ρ* with coefficients 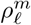, we can then apply Perseval’s theorem

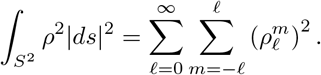

So the Willmore energy is simply the sum of squares of the SH coefficients.

This means we can interpret the power spectrum of *ρ* as the distribution of Willmore energy over each frequency *ℓ*

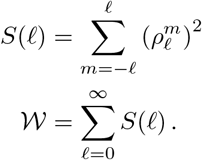

This power spectrum is rotation-invariant. [63]

#### 3.5 Implementation

In the discrete setting, a smooth surface is replaced by a mesh. This mesh consists of the sets V, E and F: the vertices, edges and faces respectively. We will only consider meshes where each face is a triangle, so that each face defines a plane and has a well-defined normal. Faces are oriented such that the normals consistently point outwards. The algorithm closely follows the methods proposed by Crane et al. [25] and Ye et al. [26, 53].

##### 3.5.1 Curvature

The integrated mean curvature is defined over the edges [64]

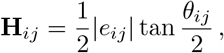

where *θ*_*ij*_ is the bending angle between two face normals *n*_*i*_ and *n*_*j*_, and *e*_*ij*_ is the edge shared by those faces (see fig. S7). The integrated mean curvature of a face is simply the sum of its edge curvatures

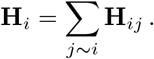

As this is an integrated quantity, it can be converted back to a pointwise quantity by dividing by the face area

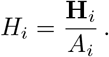

**Fig S7.**
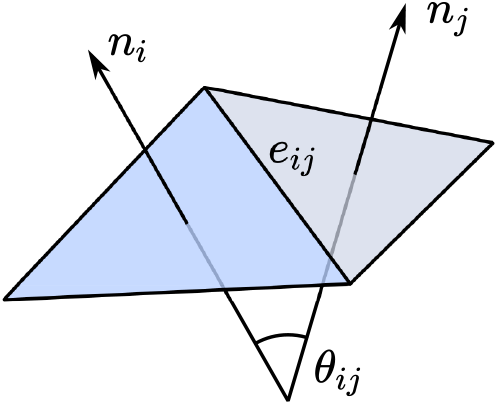
Bending angle *θ*_*ij*_ between face normals *n*_*i*_ and *n*_*j*_.

##### 3.5.2 Hyperedges and the discrete Dirac operator

We can associate with each edge the so-called *hyperedge*

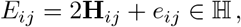

a quaternion with scalar part equal to its integrated mean curvature and imaginary part equal to its immersion in Im ℍ. Let *λ* be a function *λ* : *F* → ℍ. We then define the face-based intrinsic Dirac operator as

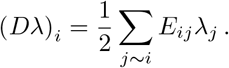

Note that this differs from the definition given by Ye et al. [26] since we drop the cosine factor. We can then solve the intrinsic Dirac equation for a function *ρ* : *F* → ℝ

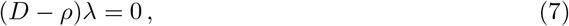

where *D* is an | *F*| × | *F*| quaternion matrix and the values of *ρ* are subtracted on the diagonal.

We then have a discrete spin transformation (*cf*. eq. (3))

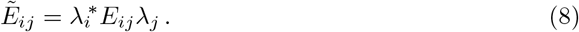

The edges 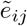 can be found as the vector (imaginary) part of 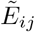.

The length element |*df* | in eq. (6) is discretized over every triangle as 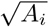

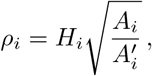

where 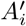 is the area of the corresponding triangle on the sphere.

##### 3.5.3 Numerical methods

Since most numerical packages can not work directly with quaternionic matrices, we represent a matrix Q ∈ ℍ ^*m*×*n*^ as a real block matrix Q^′^ ∈ ℝ ^4*m*×4*n*^. Each quaternion is replaced by a 4 × 4 block that has the same properties under ordinary matrix multiplication

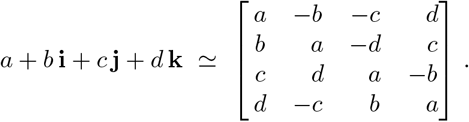

The conjugate quaternion is equivalent to the transpose of the matrix.

Solving eq. (7) directly has two problems. First, *ρ* might not correspond to a valid spin transformation at all. In that case the Dirac equation has no solution. For example, setting *ρ* = 0 everywhere on the sphere is clearly invalid, since it is impossible to flatten a sphere, removing all the curvature. So the constraint is relaxed to an eigenvalue problem [25]

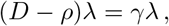

where *γ* ∈ ℝ is the smallest eigenvalue. The Dirac equation will then be satisfied where *ρ* is shifted by a small constant

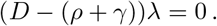

The second problem is that the solutions might not be smooth. To constrain the solution-space, it is solved in the vertices instead of the faces. A triangle mesh usually has |*F*| ≈2 |*V*|, so there are fewer degrees of freedom. If we let A be the face-to-vertex averaging matrix, we then solve the following eigenvalue problem

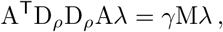

for the smallest eigenvalue *γ*. M is the diagonal vertex mass matrix. The solution is averaged back onto the faces by calculating A^T^*λ*.

We can solve this problem efficiently with the inverse power method

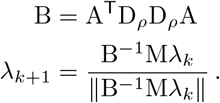

Since B is positive definite and sparse, the conjugate gradient method is an effective way to solve this system. Instead of using a random initial guess, we set *λ* to unity. This scheme converges very quickly, so we typically can stop after only three iterations.

After finding *λ*, the hyperedges are calculated by eq. (8). Integrating back to vertex positions then becomes a system of equations

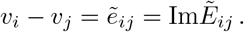

